# Complex polymorphisms in endocytosis genes suggest alpha-cyclodextrin as a treatment for breast cancer

**DOI:** 10.1101/152405

**Authors:** Knut M. Wittkowski, Christina Dadurian, Martin P. Seybold, Han Sang Kim, Ayuko Hoshino, David Lyden

## Abstract

Most breast cancer deaths are caused by metastasis and treatment options beyond radiation and cytotoxic drugs, which have severe side effects, and hormonal treatments, which are or become ineffective for many patients, are urgently needed. This study reanalyzed existing data from three genome-wide association studies (GWAS) using a novel computational biostatistics approach (muGWAS), which had been validated in studies of 600–2000 subjects in epilepsy and autism. MuGWAS jointly analyzes several neighboring single nucleotide polymorphisms while incorporating knowledge about genetics of heritable diseases into the statistical method and about GWAS into the rules for determining adaptive genome-wide significance.

Results from three independent GWAS of 1000–2000 subjects each, which were made available under the National Institute of Health’s “Up For A Challenge” (U4C) project, not only confirmed cell-cycle control and receptor/AKT signaling, but, for the first time in breast cancer GWAS, also consistently identified many genes involved in endo-/exocytosis (EEC), most of which had already been observed in functional and expression studies of breast cancer. In particular, the findings include genes that translocate (*ATP8A1, ATP8B1, ANO4, ABCA1*) and metabolize (*AGPAT3, AGPAT4, DGKQ, LPPR1*) phospholipids entering the phosphatidylinositol cycle, which controls EEC. These novel findings suggest scavenging phospholipids via alpha-cyclodextrins (αCD) as a novel intervention to control local spread of cancer, packaging of exosomes (which prepare distant microenvironment for organ-specific metastases), and endocytosis of β1 integrins (which are required for spread of metastatic phenotype and mesenchymal migration of tumor cells).

Beta-cyclodextrins (βCD) have already been shown to be effective in *in vitro* and animal studies of breast cancer, but exhibits cholesterol-related ototoxicity. The smaller αCDs also scavenges phospholipids, but cannot fit cholesterol. An *in-vitro* study presented here confirms hydroxypropyl (HP)-αCD to be twice as effective as HPβCD against migration of human cells of both receptor negative and estrogen-receptor positive breast cancer.

If the previous successful animal studies with βCDs are replicated with the safer and more effective αCDs, clinical trials of adjuvant treatment with αCDs are warranted. Ultimately, all breast cancer are expected to benefit from treatment with HPαCD, but women with triplenegative breast cancer (TNBC) will benefit most, because they have fewer treatment options and their cancer advances more aggressively.

## Introduction

Breast cancer is the most common cancer in women worldwide.^(Rojas 2016)^ In 2016, 246,660 new U.S. cases were estimated.^(Siegel 2016)^ The highly penetrant, but rare mutations in *BRCA1* and *BRCA2* point to DNA repair deficiencies as an etiological factor, but explain only 5 to 10 percent of cases. Patients with breast cancer positive for estrogen receptor (ER) or human epidermal growth factor (GF) receptor type 2 (*HER2*) initially respond well to anti-estrogen or anti-HER2 therapy, respectively, but inevitably become refractory.^(Hayashi 2015)^

As of May, 2016, the deadline for participation in the National Cancer Institutes’ “Up For A Challenge” (U4C) breast cancer challenge, 127 single nucleotide polymorphisms (SNPs) had been associated with breast cancer in women of European ancestry ^(Burdett)^ at the conventional fixed *s* = −log(*p*) = 7.3 level for genome-wide statistical significance (GWS) ^(Barsh 2012)^ (*s* is used throughout for significance). These SNPs map to 51 genes with known function; all but 16 involved in three known pathways: 27 are associated with nuclear function (DNA repair, transcription, cell-cycle control), six with receptor signaling, ion channels, and mammary gland development (KEGG pathway hsa04915) and two with AKT signaling (hsa04064).^(Kendellen 2014)^ The U4C aimed to generate novel testable biological hypotheses (80 FR 32168).

The present evaluation is based on separate analyses of three independent populations of women of European ancestry (see Subjects). Two of the populations (EPIC, PBCS) had never been analyzed individually, because their sample size was deemed insufficient for conventional statistical approaches.

Most breast cancer deaths are not due to the primary tumor, but to metastases, often in the bone, lung, liver, and brain. The genetics results submitted under the U4C implicate dysregulation and dysfunction of endo-/exocytosis (EEC), which is involved in cell migration and invasion, as well as organ targeting, and, thus, suggest overall downregulation of phosphoinositides (PI) as a novel treatment strategy against metastases. The hypothesis that alpha-cyclodextrin (αCD), which scavenges phospholipids, is effective in reducing migration of breast cancer tumor cells was subsequently confirmed in an *in vitro* study. Taken together, the results suggest (derivatives of) αCD as a potential treatment for carcinomas without the side effects of radiation and cytotoxic drugs or radiation.

## Materials and Methods

### Ethics Statement

The study was approved by The Rockefeller University IRB on Aug 24, 2015 (ref# 330390, exempt).

### Subjects

This reanalysis is based on data from three GWAS in mostly ER^−^ (including some PR^−^ and/or HER2^−^) women of European ancestry:

a. the NHS cases from the Nurses’ Health Study as part of the Cancer Genetic Markers project (CGEM, phs000147/39389-2/GRU, 1145 cases / 1142 controls),^(Hunter 2007; Haiman 2011)^
b. ER^−^ cases from the nested case-control study of the European Prospective Investigation into Cancer (EPIC, phs000812/39395-2/HMB-PU, 511 cases / 500 controls),^(Siddiq 2012)^
c. ER^−^ cases from the Polish Breast Cancer Case-Control Study (PBCS, phs000812/39397-2, 543 cases / 511 controls),^(Siddiq 2012)^

The EPIC and PBCS studies are part of the Breast and Prostate Cancer Cohort Consortium GWAS (BPC3), which was supported by the National Cancer Institute (NCI) under cooperative agreements U01-CA98233, U01-CA98710, U01-CA98216, and U01-CA98758 and the Intramural Research Program of the NCI, Division of Cancer Epidemiology and Genetics (see https://www.synapse.org/#!Synapse:syn3157598/wiki/232630 for further details).

### Statistical Methods

In this analysis, conventional single-SNP GWAS (ssGWAS) are complemented with a computational biostatistics approach (muGWAS, GWAS using muStat ^(Wittkowski 2012)^) that incorporates knowledge about genetics into the method ^(Wittkowski 2010 Sections 4.3.4 and 4.4.2; Wittkowski 2013)^ and knowledge about the nature of GWAS into the decision strategy.^(Wittkowski 2014)^

Statistical methods tend to have higher power if they are based on more realistic assumptions, which, in biology, tend to be weak. In contrast, methods based on stronger assumptions, such as additivity of allelic effects and independence of SNPs within an linkage disequilibrium (LD) block (LDB), may generate more significant results when errors happen to fulfill these assumptions than for true effects. With millions of test statistics calculated, even a rare false positive result due to model-misspecification (1/10,000 tests, say), may result in the 100 most significant results all being false positives. U-statistics for multivariate data in GWAS (muGWAS) rely only on weak, realistic assumptions, but require large amounts of memory and GPU enabled cloud instances, which became available only after 2001 and 2009, respectively.

After excluding non-informative or low-quality SNPs and SNPs in high LD with an immediate neighbor ^(Ioannidis 2009)^ (20-25%) to avoid loss of power when including irrelevant SNPs ^(Li 2012)^, an initial traditional ssGWAS was performed, using the u-test for univariate data.^(Mann 1947; Wilcoxon 1954; Kruskal 1957)^ The same data was then analyzed using a u-test for genetically structured multivariate data.^(Wittkowski 2013)^ U-statistics avoid model-misspecification biases by replacing linear/logistic^(Wu 2010b)^ with non-parametric kernels.^(Li 2012)^

Below, we describe the assumptions about genetics and GWAS that are implemented in the statistical method and decision strategy and refer to published empirical validation of this approach.

#### 1.1 Heterodominance

A particular SNP is not assumed to be either recessive (aA = aa), additive (aA = (aa+AA)/2), or dominant (aA = AA), but merely monotonic (aa < aA < AA). Accordingly, the information contributed by a particular SNP is represented as a matrix detailing for each of the *n*×*n* pairs of *n* subjects whether the genetic risk carried by the row subject is lower “<”, the same “=”, or higher “>” than the risk of column subject, or unknown (“?”) in case of missing data in one or both of the subjects. Below, the possible genetic risk constellations (left) are compared to models with different degrees of dominance (right). While the left matrix is similar to the matrix for dominant effects (all non-zero elements are ±2), the (logical) inequalities are not (numerically) equivalent. In effect, the single-SNP results based on the adaptive u-scores for aa, aA, and AA are similar to results from the Cochran-Armitage test for additive co-dominance, ^(Cochran 1954; Armitage 1955)^ which uses fixed scores 0, 1, and 2.

| X~Y | aa | aA | AA | ?? | X~Y | aa | aA | AA | ?? |
| --- | --- | --- | --- | --- | --- | --- | --- | --- | --- |
| aa | = | < | < | ? | aa | $\pm 0$ | $+2 / +1 / \pm 0$ | $+2$ | ? |
| aA | > | = | < | ? | aA | $\pm 0 / -1 / -2$ | $\pm 0$ | $+2 / +1 / \pm 0$ | ? |
| AA | > | > | = | ? | AA | $-2$ | $\pm 0 / -1 / -2$ | $\pm 0$ | ? |
| ?? | ? | ? | ? | ? | ?? | ? | ? | ? | ? |

#### 1.2 LD-structure

A basic assumption underlying GWAS, in general, is that a disease locus should be in LD with both neighboring SNPs (unless they are separated by a recombination hotspot). Hence, the information from two neighboring SNPs is not numerically ADD-ed, but logically AND-ed using the function ∧

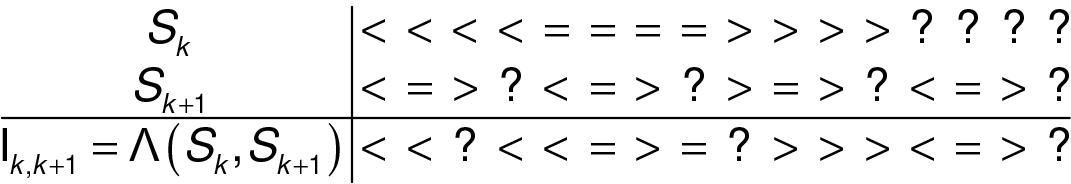

As muStat allows variables to be correlated, other SNPs within an LDB may be in LD, too, yet there is no formal representation of more distant LD. Non-informative SNPs added between LDBs prevent intervals from spanning LDBs.

#### 1.3 Cis-epistasis, including compound-heterozygosity

To account for interactions between functional polymorphisms,^(Aslibekyan 2013a)^ a natural extension of ∧ is then used to combine information from corresponding elements of the *n*×*n* matrices containing information about neighboring pairs. Assuming, without loss of generality, the case of only four SNPs within in the same LDB, the aggregated diplotype information for one pair of subjects is

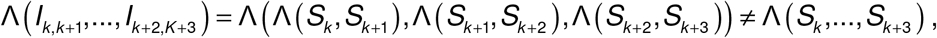

which can be one of the following (invariant to permutaitons π):

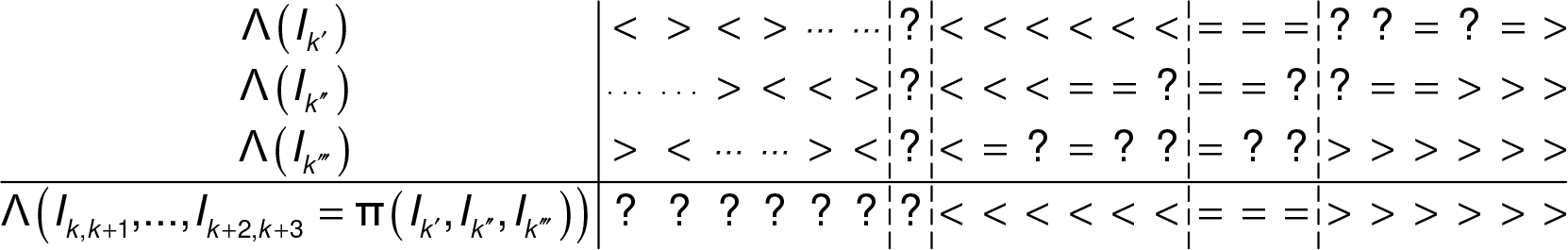

From the above inequality, the results typically differ when SNPs from the same tag sets appear in different permutations, which increases the resolution over methods assuming commutativity.

#### 1.4 Test statistic

From the resulting *n*×*n* matrix *W* (say), one calculates each subject’s risk score *u_i_* (-*n* < *u_i_* < *n*) as the number of subjects having lower risk, minus the number of subjects having higher risk, i.e., #(*W_ij_* = “<”)_*j*_ - #(*W_ij_* = “>”)_*j*_. These scores are then compared between cases and controls using a standard linear score test.^(Hajek 1967)^

#### 1.5 Regularization

Since it is unknown *a priori*, whether a minor allele is dangerous, irrelevant, or protective, all combinations of (−1,0,+1) “polarities” are applied to the SNPs *S_k_*, …, *S*_*K*+3_, resulting in many highly dependent test statistics being calculated for the diplotypes surrounding a given SNP. The test statistic chosen is the one that has the highest u(-log(*p*), *IC*) score, where the information content (IC) is the proportion of pairwise orderings in *W* that can be decided (≠ “?”) for a given choice of polarities. This approach avoids over-fitting (highly significant results based on a small subset of unusual subjects) without the need to choose arbitrary regularization cut-offs.^(Frommlet 2016)^

#### 2.1. Adaptive genome-wide significance

The traditional p-value cut-off of *s* = 7.3 for GWS has been widely criticized as overly conservative,^(Pearson 2008; Panagiotou 2012)^ yet few alternatives have been formally derived. Here, we replace a fixed cut-off for GWS with an empirical ^(Aslibekyan 2013a)^ adaptive (study-specific) cut-off (aGWS) that automatically accounts for the specifics of the population studied, the chip used, differences in minor allele frequency (MAF,) and GWAS being non-randomized.^(Wittkowski 2014)^ As previously discussed,^(Wittkowski 2014)^ the expected distribution in a ssGWAS QR plot is a mixture of univariate distributions whose carriers vary by MAF, because the most significant result possible depends on MAF when outcomes are bounded (allele counts 0, 1, 2). Hence, it is a convex curve, rather than a straight line;^(Wittkowski 2014)^ see, for instance, CGEM chromosomes 14-17, 19, and 22 (S1 Fig 2). In a whole genome (WG) plot, this curvature may not be apparent (see Fig 1, below), when some chromosomes’ QR curves are concave because of true association, which is expected in a familial disease or with systematic unrelated differences between non-randomized populations. Hence, an apparently straight line in a WG plot may be due to concave curves in chromosomes with true positives and convex curves in others canceling each other out. With muGWAS, where many dependent tests are performed at overlapping window positions, the expected QR curve (see S1 Fig 3) may be even more convex. The expected distribution curve is estimated from the 50% of chromosomes with the fewest outliers rising above a convex fit.^(Wittkowski 2014)^ The empirical adaptive (study-specific) aGWS cutoff is the median apex (highest point) of a convex curve fitted against these chromosomes’ QR plot.

#### 2.2. Replication

Complex diseases may involve different SNPs in high LD with causal variants across populations,^(Pickrell 2016)^ epistasis between several SNPs per locus, several loci per gene, and several genes per function, with risk factors differing across populations (see above). Hence, we will consider SNPs within a locus, loci within a gene, and genes with a direct mechanistic relationship (paralogs, binding partners, …) for replication. ^(Peng 2010; Aslibekyan 2013a)^ Results are considered “replicated” if supportive results are significant at the aGWS/2 level.

**Validation**: The above approaches have been validated in two published analyses, where previous analyses using ssGWAS and fixed GWS also had identified not more than a few apparently unrelated SNPs.

- In epilepsy,^(Wittkowski 2013)^ muGWAS confirmed the Ras pathway and known drug targets (ion channels, *IL1B*). In that analysis, muGWAS was also compared with a parametric analogue, logistic regression with interaction terms for neighboring SNPs (lrGWAS). muGWAS produced fewer apparent false positives (isolated highly significant results far away from coding regions) ^(Wittkowski 2013 Suppl. Fig 2)^ and higher sensitivity for genes downstream of Ras, which are involved in more complex cis-epistatic interactions,^(Wittkowski 2013, Fig 3, blue)^ than ion channels, which were also implicated by lrGWAS.^(Wittkowski 2013, Fig 3, red)^
- In autism,^(Wittkowski 2014)^ muGWAS identified sets of mechanistically related genetic risk factors for mutism in autism (independently confirmed in functional studies ^(Guglielmi 2015)^ and a pathway network analysis ^(Wen 2016)^). In, ^(Wittkowski 2014)^, adaptive GWS was validated against three analyses with randomly permutated phenotypes. Only one gene (DMD, not aGWS) appeared in one of the other analyses (also not aGWS). Moreover, there is no noticeable overlap between aGWS genes between breast cancer and either mutism^(Wittkowski 2014)^ or epilepsy ^(Wittkowski 2014, Suppl. Fig 7)^, while there is considerable functional overlap between mutism in autism and epilepsy, as expected.

### *In vitro* Assay

A 24-well plate (CBA-120, Cell BioLabs Inc., San Diego, CA) with CytoSelect Wound Healing Inserts was warmed up at room temperature for 10 min. A cell suspension used contained 0.5-1.0 × 10^6^ cells/ml in media containing 10% fetal bovine serum (FBS) was prepared and 1 mL of this suspension was added to each well. Cells were then incubated for 12 h, after which time the insert was removed and cells were washed with new media to remove dead cells and debris. FBS with/without CDs (Sigma-Aldridge, St. Louis, MO) was added to start the wound healing process. Cells were incubated for 2 h, washed with PBS, fresh control media was added, and cells were incubated for another 12 h. After removing the fixation solution, 400 μL of Cell Stain Solution were added to each well and incubated for 15 min at room temperature, after which stained wells were washed thrice with deionized water and left to dry at room temperature. Cells that migrated into the wounded area or protruded from the border of the wound were visualized and photographed under an inverted microscope to determine migrated cell surface area. https://www.cellbiolabs.com/sites/default/files/CBA-120-wound-healing-assay.pdf

## Results

### Additional ssGWAS CGEM results complement known breast cancer risk factors

The original CGEM analysis had identified two SNPs (rs1219648: *s* = 5.49, rs2420946: 5.46) in the fibroblast GF receptor *FGFR2* ^Entrez Gene 2263 (Hunter 2007)^ which affects mammary epithelial cell growth and migration,^(Czaplinska 2014)^ and a SNP (rs10510126: 6.25, >1 MB apart from *FGFR2*) which was subsequently located to a long variant of the mitotic checkpoint protein *BUB3* ^9184^.

**Fig 1:**
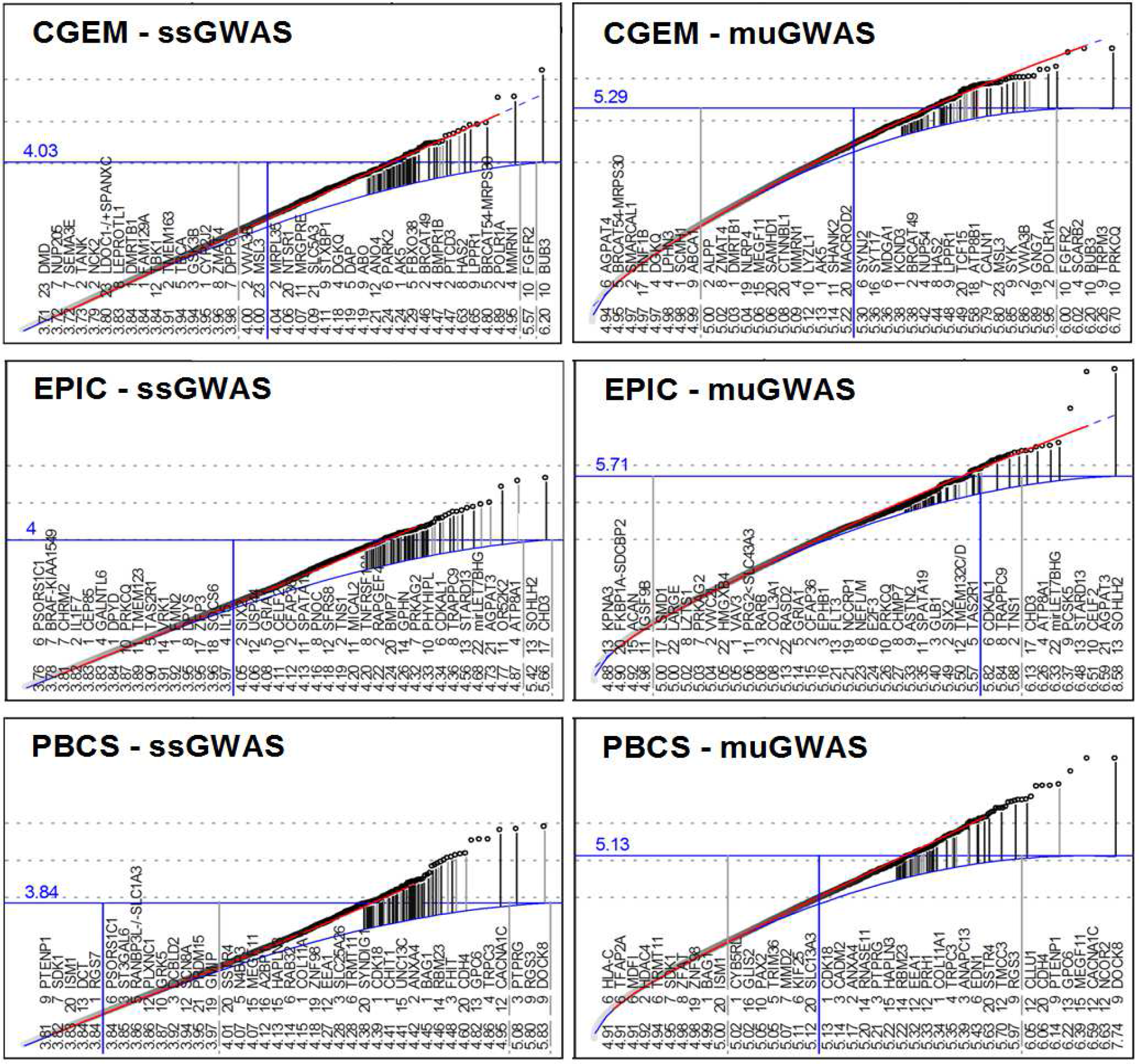
GWAS Quantile-Rank (QR) plots. Left: ssGWAS, right: muGWAS (each point represents the most significant result among all diplotypes centered at the same SNP) Results are ranked by significance (bottom). For the most significant results and other results of interest, the location of SNPs to genes is shown in S1 Fig 5. Upper curve (red): convex fit against points; dashed extension: projection; lower curve (blue): population-specific expectation. Vertical lines between curves connect the highest *s*-values (-log10 *p*) of a gene (dot) with its expected value for genes with known function. Light gray vertical lines indicate genes omitted from the list because of low reliability (either low μlC or reliance on a single SNP), respectively. Genes to the right of the vertical dark line are above the aGWS cut-off. See S1 Fig 1 for Manhattan plots. The horizontal solid line at highest point at the end of the expected curve indicates the estimate for adjusted GWS (aGWS). All points above the horizontal line (and genes to the right of the vertical blue line) are “significant” at the aGWS level.

These two genes are also the only genes in the present analysis with SNPs above the diagonal in the summary ssGWAS quantile-rank (QR, often: QQ) plot (Fig 1 left), although the QR plots of several individual chromosomes show association in chromosomes 4 (the *SNCA-MMRN1* ^22915^ region), 5 (breast cancer associated transcript *BRCAT54* ^100506674^, non-coding), 6 (*PARK2*^5071^, the Parkinson’s disease [PD] ubiquitin ligase Parkin), and 9 (*LPPR1* ^54886^, phospholipid phosphatase-related 1) (S1 Fig 2).

In the present analysis, a total of 22 genes and *BRCAT49*^(Iyer 2015)^ reached aGWS in CGEM (Fig 1, left, blue). A total of 21, 11, and 24 genes with known function or relation to breast cancer exceeded muGWAS aGWS in CGEM, EPIC, and PBCS, respectively.

### Novel ssGWAS aGWS results in EPIC and PBCS complement CGEM results

In EPIC, the two most significant SNPs (rs4791889: 5.66 and rs9596958: 5.42) are located 4.5 kB upstream of the chromodomain helicase DNA binding protein *CHD3* ^1107^ and the transcription factor (TF) *SOHLH2* ^54937^, respectively (see S1 Table 2 and S1 Fig 5).

In PBCS, the two most significant SNPs (rs2297075: 5.83, rs943628: 5.55, 100 kB apart) are located in *DOCK8* ^81704^, a guanine nucleotide exchange factor for Rac1, which drives mesenchymal cell movement.^(Wang 2015b)^ Significance of *FGFR2* relies on the two previously reported and a third SNP (rs11200014) within intron 2.^(Cui 2016)^ Significance in *BUB3* is driven by three SNPs in high LD (rs10510126, rs17663978, rs7916600, spanning 30kB). These findings are consistent with significance of the top five SNPs in ssGWAS depending on a single polymorphism each. Lack of evidence in EPIC and PBCS (S1 Table 2) is consistent with different variations developing in divergent European populations.

### muGWAS aGWS results are cross-validated across CGEM, EPIC, and PBCS

In CGEM, the top gene was the phospholipid/diacylglycerol (DAG)-dependent protein kinase *PRKCQ*^5588^ (chr10: 6,540,724-6,573,883), which induces cell migration and invasion.^(Belguise 2012; Zafar 2014)^ The same SNP (rs661891) was also implicated in EPIC. The three most significant SNPs and the most significant regions in muGWAS were all located within the same 34 kB LDB. The second most significant gene was a long EST of the transient receptor potential cation channel *TRPM3* ^80036^, which controls oncogenic autophagy in renal cell carcinoma,^(Hall 2014)^ supported by a part of the promoter region of the shorter main form in PBCS. *BUB3* was also significant in muGWAS, followed by the endo-/lysosomal receptor *SCARB2* ^950^ and the nuclear RNA polymerase subunit *POLR1A* ^25885^ (rs10779967).

In EPIC, the top gene in muGWAS (as in ssGWAS), was the TF *SOHLH2*, followed by *AGPAT3* ^56894^ (rs8132053 in CGEM and EPIC), whose paralog *AGPAT4* ^56895^ is included in Fig 1 (4.94, right panel, CGEM). *CELF2*^10659^, an RNA binding protein, and *STARD13* ^10948^, a breast cancer tumor suppressor that regulates cell migration and invasion ^(Hanna 2014)^ also reached aGWS. *CHD*^364663^ depends entirely on SNP rs4791889 (see Statistical Methods, 2.2. Replication, for replication criteria).

In PBCS, the top gene in muGWAS, as in ssGWAS, was *DOCK8* ^81704^, followed by the nuclear receptor corepressor *NCOR2* ^9612^, which has been implicated in tamoxifen resistance in breast cancer.^(van Agthoven 2009; Zhang 2013b)^ *CACNA1C* ^775^ (3^rd^) is highly up-regulated in breast cancer.^(Wang 2015a)^ The multiple epidermal GF-like domains protein 11 (*MEGF11* ^84465^, 4^th^), like *MEGF10* ^84466^ an ortholog of *C. elegans* Ced-1 and the *Drosophila* draper, had been implicated in colorectal cancer. ^(Cicek 2012)^

Both CGEM and EPIC have a significant P-type ATPase, which import phosphatidylserine (PS, *ATP8B1* ^5205^) and phosphatidylcholine (PC, *ATP8A1* ^10396^), respectively, the substrates for phos-pholipase D (PLD) to produce phosphatidic acid (PA) for the synthesis of phosphatidylinositol (PI).^(Daleke 2007)^ *BMR7*^655^ (ss: 4.24) and its receptor *BMRR1B* ^658^ (ss: 4.47) are significant in EPIC and CGEM, respectively, and BMP signaling is known to regulates mitotic checkpoint protein levels in human breast cancer cells, including levels of *BUB3* (see above).^(Yan 2012)^ *DGKQ* ^1609^ (rs2290405) which converts DAG into PA, was replicated in CGEM and PBCS, while *LPPR1* ^6^, which is involved in the conversion of PA into PI was replicated in CGEM and EPIC.

As expected in samples from the general population, the known risk factors for rare early-onset breast cancer (*BRCA1/2* ^672/675^, *HER2* ^2064^, *RB1* ^5925^) do not show association and many receptor-related genes are absent in ER^−^ populations. Except for the genes with highest significance in ssGWAS (*BUB3* in CGEM, *SOHL2* in EPIC, and *DOCK8* in PBCS), all of the aGWS genes in muGWAS have support in least one of the other two populations (2^nd^ block of S1 Table 2). This observation is consistent with muGWAS identifying primarily old cis-epistatic variations, rather than *de novo* mutations favored by ssGWAS. S1 Table 2 gives an overview about the significance and replication of the genes identified and supportive evidence in the literature.

### muGWAS results confirm known disease pathways in breast cancer

Consistent with the published results in the NHGRI-EBI catalog, a total of 16, 15, and 18 genes above aGWS in CGEM, EPIC, and PBCS, respectively, are involved in the three known disease pathways, such as membrane-associated receptor signaling (G protein-coupled receptors [GPCR], Fc receptors [FcR], hemagglutinin [HA], receptor tyrosine kinases [RTK], or ion channels), MAP kinases, and in nuclear proteins involved in cell cycle control, transcription, or splicing in breast cancer (Table 1).

### muGWAS results highlight Endo-/Exocytosis (EEC) as a pathway in breast cancer

The cell’s major fibronectin-binding integrin (α5β1) is key to the survival and migration of tumor cells.^(Dozynkiewicz 2012)^ Results of various expression and functional studies have pointed to EEC of β1 integrins as a functional component of “derailed endocytosis” in cancers, including breast cancer (Fig 2).^(Mosesson 2008; Morgan 2009; De Franceschi 2015)^

Among the 15 GWS genes not associated with known pathways in the NHGRI-EBI catalog (excluding the ambiguous locus between *MDM4* ^4194^ and *PIK3C2B* ^5287^), only four are involved in EEC (*PDE4D, SNX32, STXBP4, DNAJC1*, marked with “*” in S1 Table 1), all from ssGWAS of a combined analysis of nine studies,^(Michailidou 2013)^ which includet the three studies analyzed separately here. A String(^http://string-dborg/^) pathway analysis of the subset of aGWS genes that are not part of the above three pathways identified two clusters related to EEC:

- **EEC Function**: *PARK2, PTEN (from PTENP1), SYNJ2, STXBP1, UNC13* (consistent with previous functional studies, see S1 Table 3)
- **EEC Regulation**: *AGPAT3* and *DGKQ* (S1 Fig 4).

### muGWAS identified genes causing dysfunction of EEC, a known BC risk factor

Further String subset analyses and a literature review by the authors identified additional aGWS genes as related to EEC-related KEGG pathways (genes in parenthesis replaced by a related gene with known function in String). They include endocytosis (hsa04144): *DNM1* (from *MEGF11*), *EEA1, PDE4D, SNX32*, NEDD4(from *N4BP3*) (FDR = .018) and synaptic vesicle cycle (hsa04721): *STXBP1, UNC13C, VAMP2*; (FDR = .0001).

Fig 4 integrates the genes identified in the present GWAS analysis (pink, see S1 Table 1 for details) with results from expression and functional studies of β1 integrin EEC in breast cancer (see S1 Table 3 for details).

**Table 1:**
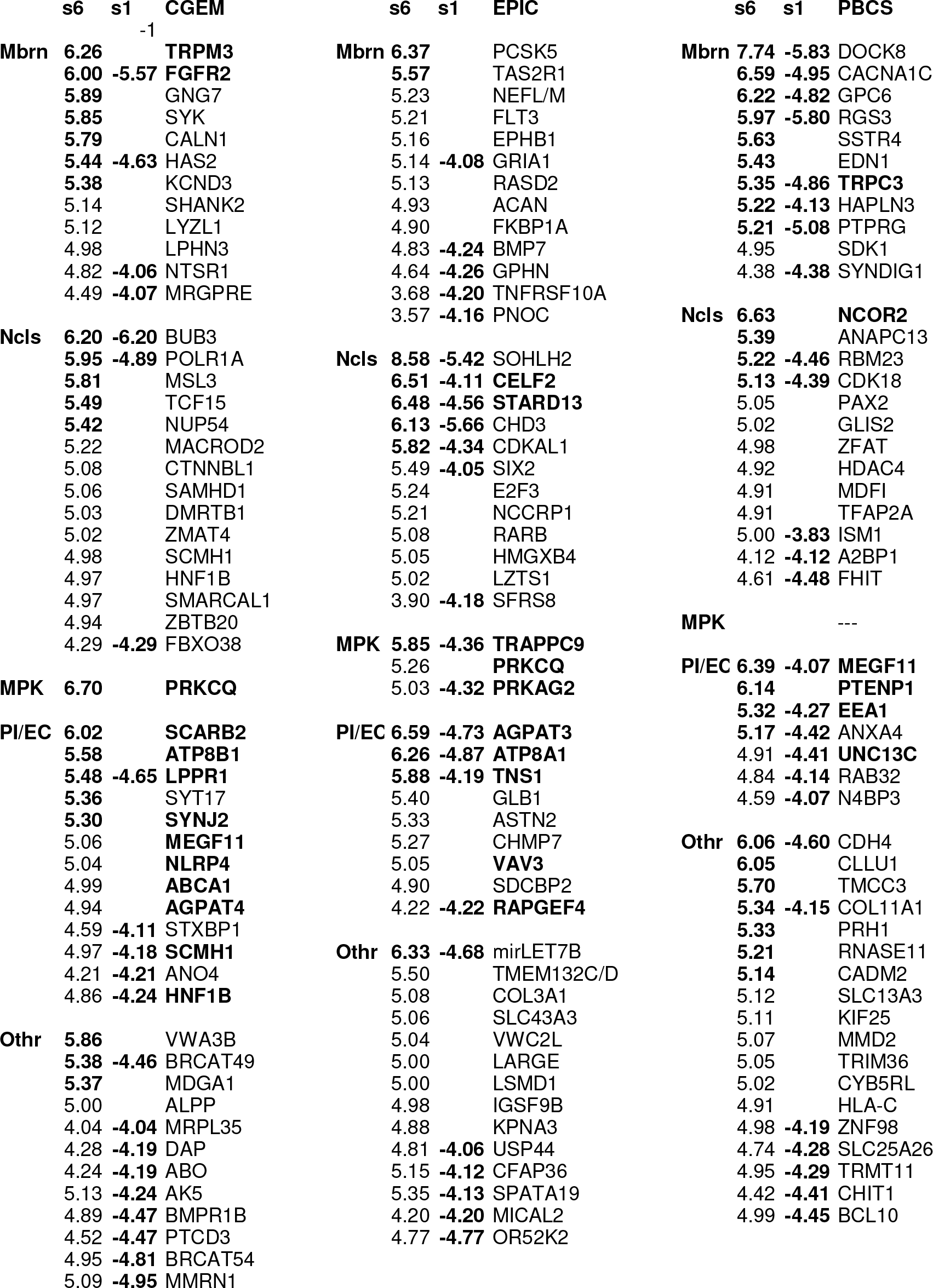
Breast cancer genes associated with pathways by Study. Within each study (major columns), genes are grouped by function. Mbrn: membrane-associated (GPCR, FcR, HA, RTK, Ion channels), Ncls: nuclear (cell cycle control, transcription, splicing), MPK: MAP kinases, PI/EC: PI cycle/EEC,.Othr: other. Within each block, muGWAS genes (Fig 1) are sorted from top by s-value (s6). s-values above aGWS (CGEM: 5.29, EPIC: 5.71, PBCS: 5.13) are shown in bold. Genes above aGWS in ssGWAS only (CGEM: 4.03, EPIC: 4.00, PBCS: 3.84) are sorted from bottom up (s1); ssGWAS results for genes also implicated in muGWAS are shown next to the muGWAS results. See S1 Table 1 for Entrez Gene identifiers and S1 Table 2 for replication across populations, which is indicated in bold names.

**Fig 2:**
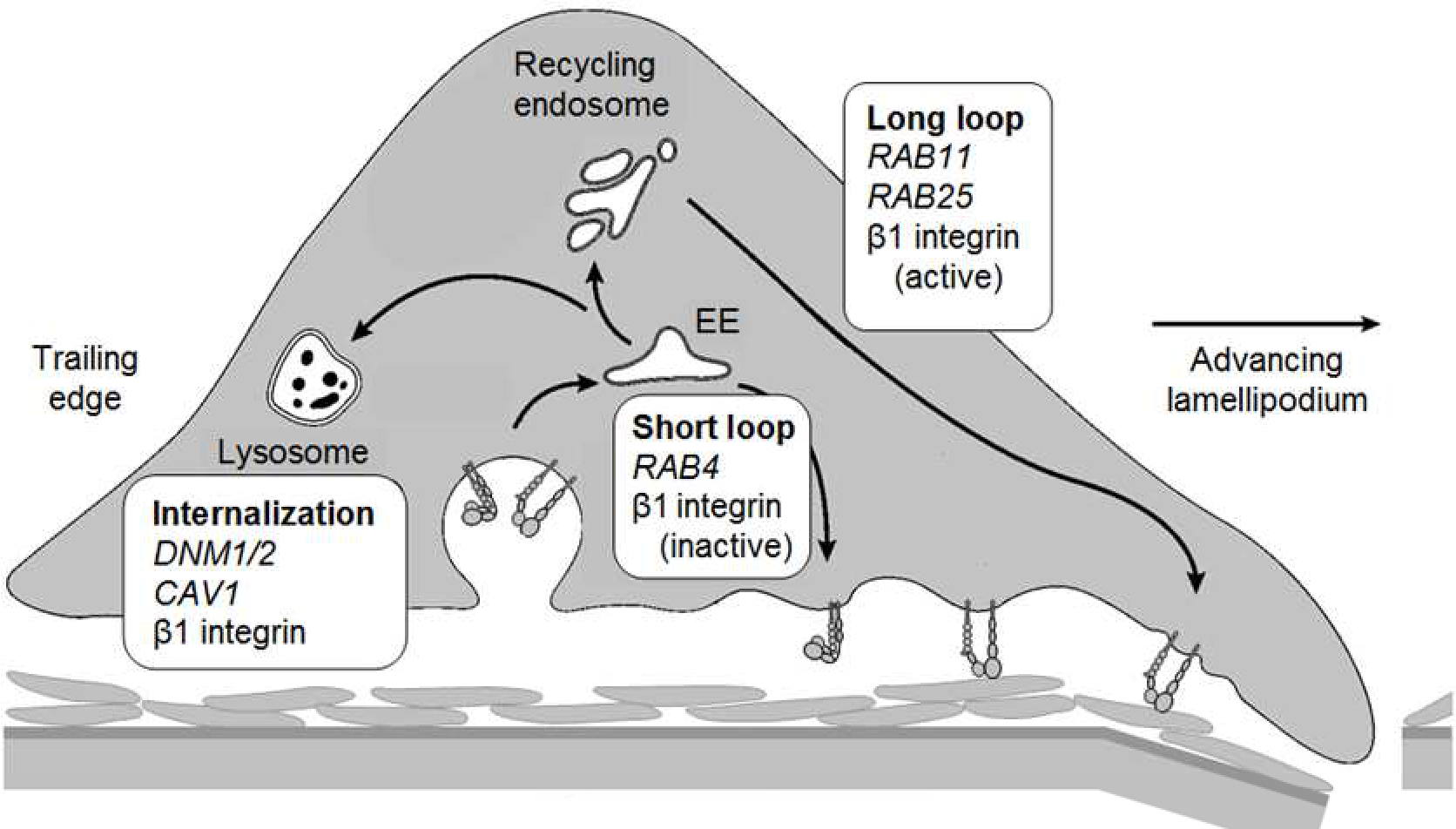
EEC of β1 Integrin underlying mesenchymal tumor cell migration and invasion. Cell migration necessitates trafficking of β1 integrin, whose internalization is controlled by dynamin. Both clathrin-and caveolin 1 (CAV1)- coated domains of the plasma membrane are involved. Once in early endosomes (EE), integrins may be sorted for degradation in lysosomes, recycled to the plasma membrane through a RAB4-dependent route, or transported to the recycling endosome (RE). Recycling from the RE requires Rab11 family members, such as RAB25 which is often aberrantly expressed in human tumors, including luminal B breast cancer.^(Mitra 2016)^ (adopted from ^(Mosesson 2008; Morgan 2009; De Franceschi 2015)^)

### muGWAS identifies PI cycle dysregulation as novel breast cancer risk factor

In relation to EEC regulation, both CGEM and EPIC identified a phospholipid-translocating ATPase, *ATP8B1* (PE) and *ATP8A1* (PS), respectively. *AGPAT3* is the second most significant gene in EPIC (mu: 6.59, ss: 4.73); *AGPAT4* is among the supportive genes in CGEM (Fig 1, mu: 4.94). Both acyltransferases transform LPA into PA. CGEM also identified the scramblase *ANO4* ^121601^ (ss: 4.21), a PS exporter, and the plasma membrane PC/PS efflux pump *ABCA1* ^19^ (mu: 4.99). For (*ATP8A1, ATP8B1, ANO4, ABCA1*), String identified functional enrichment in

GO:0097035 (biol. process) Regulation of membrane lipid distribution: FDR = 0.012
GO:0015914 (biol. process) phospholipid transport: 0.0407
GO:0005548 (mol. function) phospholipid transporter activity: 0.00968

As shown in Fig 4 (upper left corner), 8 (including 6 aGWS) genes are involved in providing the PI cycle with its substrate, PI (and the MAPK signaling pathway with PA.^(hsa04072)^.

### Results for EEC regulation and function are consistent cross populations

All three populations show aGWS association with EEC genes (CGEM: 4 in ssGWAS only / 4 in muGWAS only / one in both; EPIC: 1/0/3; PBCS: 3/1/3). Most are validated in at least one of the other two populations, either by the same SNP involved (*AGPAT3, DGKQ*), the same region (*SYNJ2, PDE4D*), the same gene (see S1 Fig 5), or a functionally related gene (*AGPAT3/AGPAT4, LPPR1/DGKQ, ATP8A1/ATP8B1, STXBP1/UNC13C, TNS1/PTENP1*, see S1 Table 2 for details).

**Fig 3:**
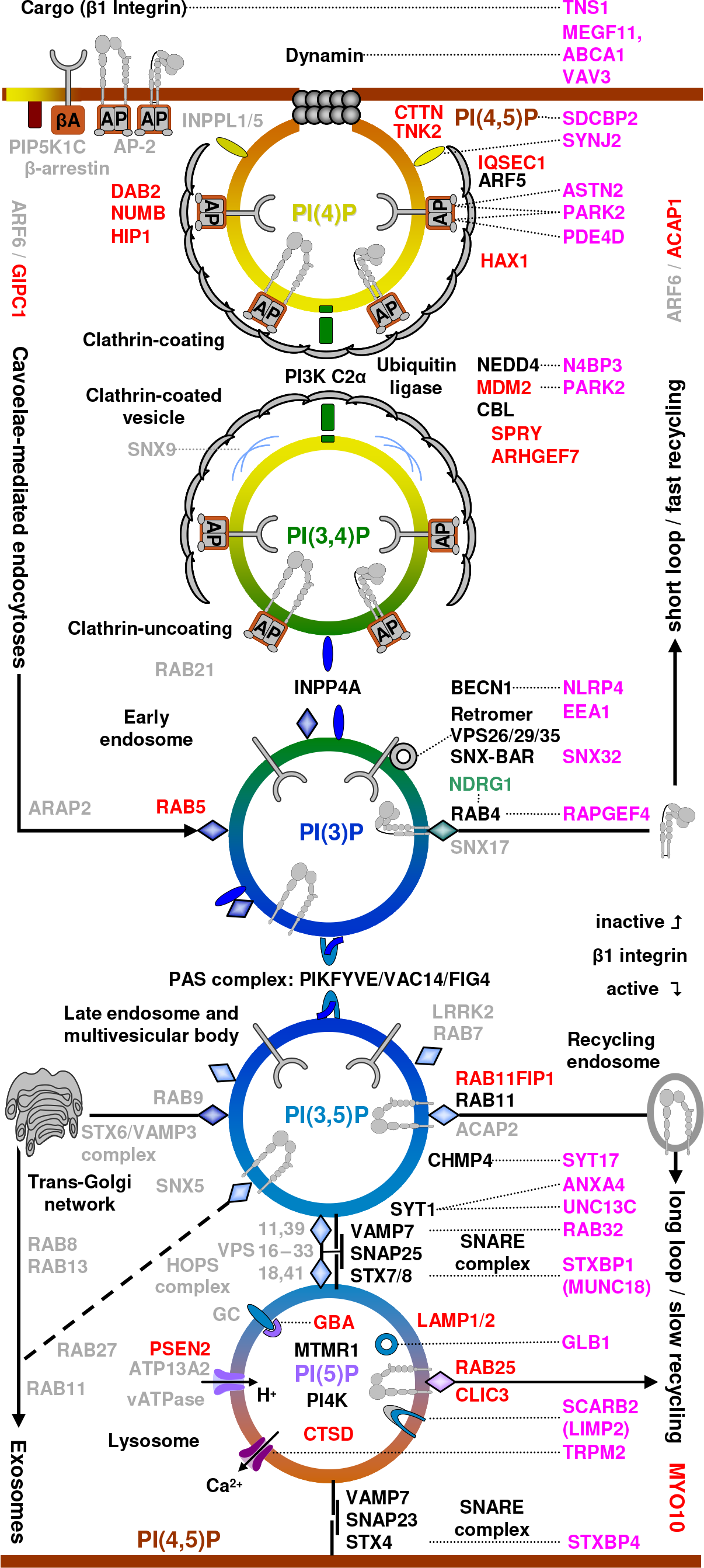
Endo-/exocytosis pathway. Pink: genes identified in this analysis, most of which have been implicated in breast cancer previously (S1 Table 1), by stage of EEC: Formation of clathrin-coated vesicles, E3 ubiquitination, separation of inactive integrin (fast recycling) from active integrins (slow recycling), sorting between secretory, lysosomal, and (slow) recycling pathway, and lysosomal degradation. Red and green genes are known breast cancer promoters and suppressors, respectively (S1 Table 3). Clathrin-mediated endocytosis (CME) begins with co-assembly of the hetero-tetrameric adaptor complex AP-2 with clathrin at PI(4,5)P_2_-rich plasma membrane sites. AP-2 in its open conformation recruits clathrin and additional endocytic proteins, many of which also bind to PI(4,5)P_2_. Maturation of the clathrin-coated pit (CCP) may be accompanied by SHIP-2-mediated dephosphorylation of PI(4,5)P_2_ to PI(4)P. Synthesis of PI(3,4)P_2_ is required for assembly of the PX-BAR domain protein SNX9 at constricting CCPs and may occur in parallel with PI(4,5)P_2_ hydrolysis to PI(4)P via synaptojanin, thereby facilitating auxilin-dependent vesicle uncoating by the clathrin-dependent recruitment and activation of PI3KC2α, a class II PI3-kinase. PI(3,4)P_2_ may finally be converted to PI(3)P en route to endosomes by the 4-phosphatases *INPP4A/B*, effectors of the endosomal GTPase Rab5. Adapted from ^(Posor 2015)^ In the early endosome, β1 integrins are sorted. Inactive integrins undergo fast “short loop” recycling; active integrins go to the late endosome / multivesicular body for slow “long group” recycling (*RAB11*), lysosomal degeneration (unless rescued by *RAB25/CLIC3*), or secretion via the trans-Golgi-network mediated by *RAB9*. Fast recycling of epidermal GF receptor drives proliferation,^(Tomas 2014)^ so one would expect gain-of-function mutations in the upper part of the Figure. Lysosomal and synaptic vesicle exocytosis share many similarities. Endolysosome-localized PIs may regulate lysosomal trafficking in early onset lysosomal storage diseases.^(Samie 2014)^ and, particularly in ageing, insufficient lysosomal degradation contributes to Alzheimer’s disease (*PSEN1, PSEN2, GLB1, CTSD*), Parkinson’s disease (*ATP13A2, GBA*),^(Colacurcio 2016)^ atherosclerosis (vATPase),^(Jaishy 2016; Chistiakov 2017)^ and type 2 diabetes (*GLB1, HEXA*).^(Tiribuzi 2011)^ (derived, in part, from KEGG pathways hsa04144, hsa04721, hsa00531, and hsa04142).

**Fig 4:**
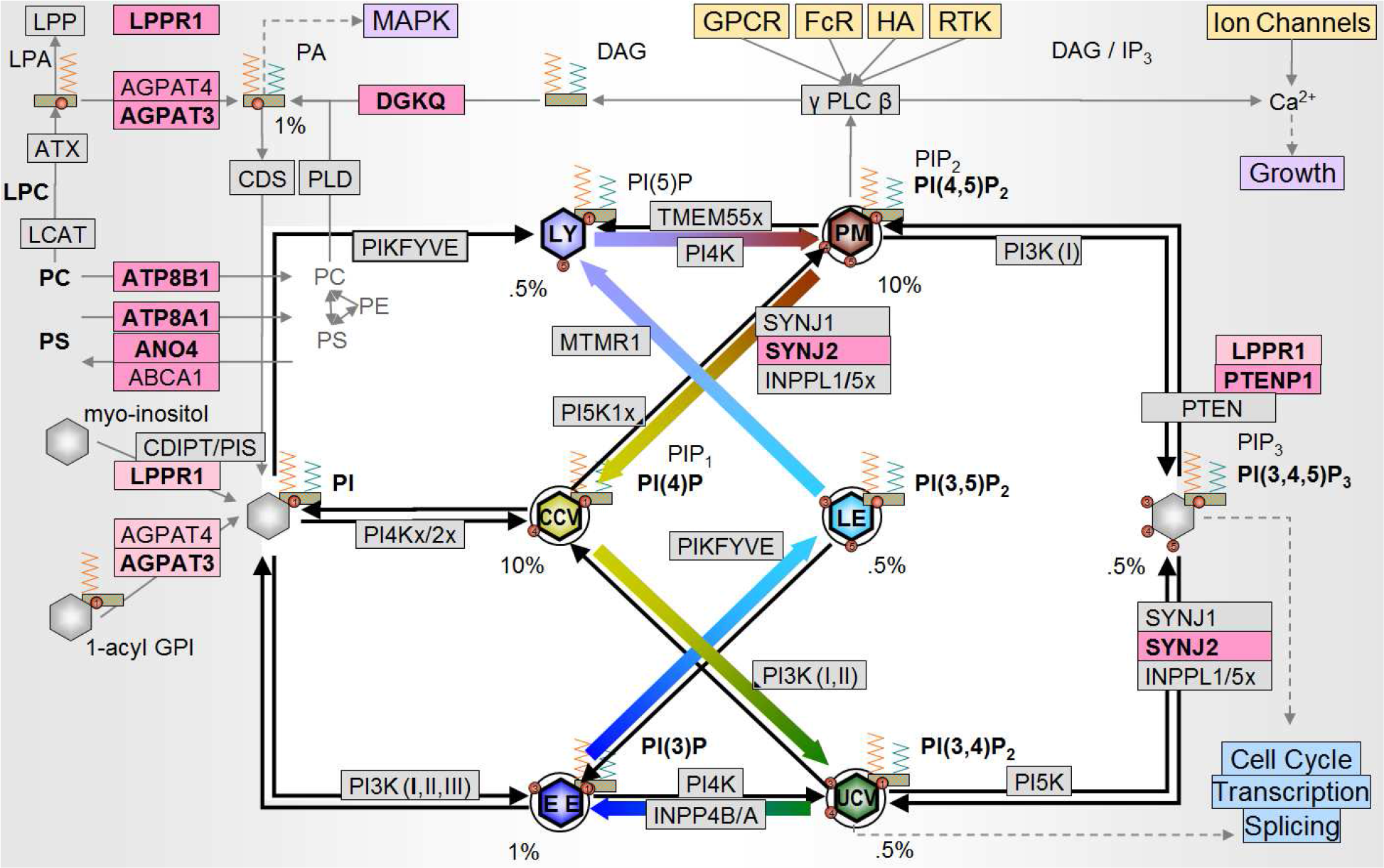
Functional relation of the PI/EC genes. PI is synthesized from myo-inositol (imported by HMIT) and PA (via CDP-DAG) which can be synthesized from lysophosphatic acid (LPA), PC, or PS, or salvaged from IP_3_ and DAG. It can also be synthesized from 1-acyl GPI. Arrows: PIs are phosphorylated at a 3-, 4-, or 5- position by Pl-kinases (left to right) and hydrolyzed by phosphatases (right-to-left). Genes associated with breast cancer in this GWAS are highlighted in pink (bold: aGWS). See Table 1 for other box colors. Colored arrows in the center indicate the sequence of PIs involved in EEC (Fig 3). Percent values indicate approximate proportion of phospholipids.^(Viaud 2016)^.

### PI supply into the PI cycle as a drug target in breast cancer

After loss-of-function in *PTEN* and gain-of-function in *PI3K* suggested a mechanism for upregulation of PI(3,4,5)P_3_ in cancer, blocking *PI3K* with Wortmannin ^(Powis 1995)^ or related drugs ^(McNamara 2011)^ were considered for treatment of cancers, including breast cancer. Upregulation in PI(3,4)P_2_ (gain-of-function in *SYNJ1/2* or *INPPL1*^(Bunney 2010)^) and PI(3)P (gain-of-function in *INPP4B*),^(Woolley 2015)^ have also been associated with breast cancer. Recently, components to lower PI(3,4)P_2_ by inhibiting *SYNJ2* have been identified.^(Ben-Chetrit 2015)^

Targeting individual phosphotransferases is unlikely to succeed given the robustness of the PI cycle.^(Powis 1995)^ All PIs regulating EEC, except for the evolutionarily recent *MTMR1* link (Fig 4), are regulated by both three kinases and three groups of phosphatases. Given the plethora of genes involved in EEC (Fig 3) identifying the appropriate set of phosphotransferase for a given patient to interfere with endocytosis or to correct for functional deficits in exocytosis may be impractical.

Regulating EEC by controlling the availability of phospholipids, however, while leaving functional interactions within the PI cycle intact, may be feasible. In fact, adding of either exogenous PS or PE led to an enhancement of endocytosis.^(Farge 1999)^ As EEC is an essential and highly conserved mechanism for tissue morphogenesis ^(Emery 2006; Bokel 2014)^ and neuronal migration,^(Wilson 2010; Cosker 2014; Kawauchi 2015)^ loss-of-function mutations would likely terminate embryonal development. Accordingly, the overall effect of the variations identified (S1 Table 3) is likely gain-of-function.

### HPaCD is more effective than HPbCD against migration of breast cancer cells

In 2014, it was reported that the benefit attributed to the neurosteroid allopregnanolone in the treatment of Niemann-Pick type C (NPC) disease was due to the expedient, 2-hydroxypropyl-beta-cyclodextrin (HPβCD). Cyclodextrins are hydrophilic rings of ≥6 starch molecules (Fig 5). The lipophilic cavity can transport lipid drugs, such as allopregnanolone. Empty CDs, at therapeutic doses, form a pool in the aqueous phase into which, in the case of βCDs, cellular cholesterol is extracted,^(Ohtani 1989)^ the mechanism of action in NPC.^(Vance 2014)^

**Fig 5:**
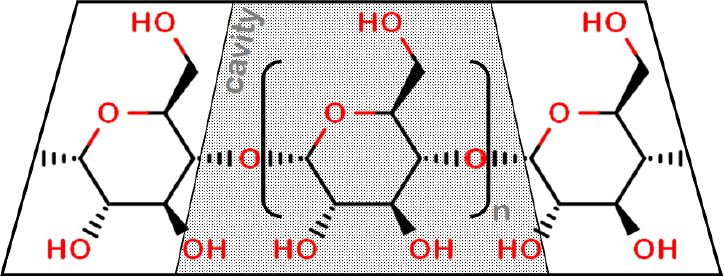
Structure of cyclodextrins. Cyclodextrins are toroids formed of six (n=4, αCD), seven (n=5, βCD), or eight (n=6, γCD) starch molecules. The cavity is lipophilic, while the surface is hydrophilic.

Given the focus on cholesterol in NPC, it has often been overlooked that βCDs also scavenge phospholipids. The above GWAS results (Table 1) suggested defects in phospholipid, rather than cholesterol function. Hence, the efficacy of HPβCD in breast cancer might be due to its ability to scavenge phospholipids.

HPβCD is known to inhibit migration of human MDA-MB 231 breast cancer cells.^(Liu 2007; Donatello 2012; Guerra 2016b, Figure 3B)^ To determine whether inhibition of migration is caused by HPβCD depleting cholesterol, as assumed previously, or by it depleting phospholipids, as implicated by the novel genetics results, the published activity from wound healing experiments comparing HPβCD against control was replicated, and complemented with novel activity results comparing HPαCD against control, both in MDA-MB 231 (ER^−^) and MCF-7 (ER^+^) human breast epithelial cell lines.

**Fig 6:**
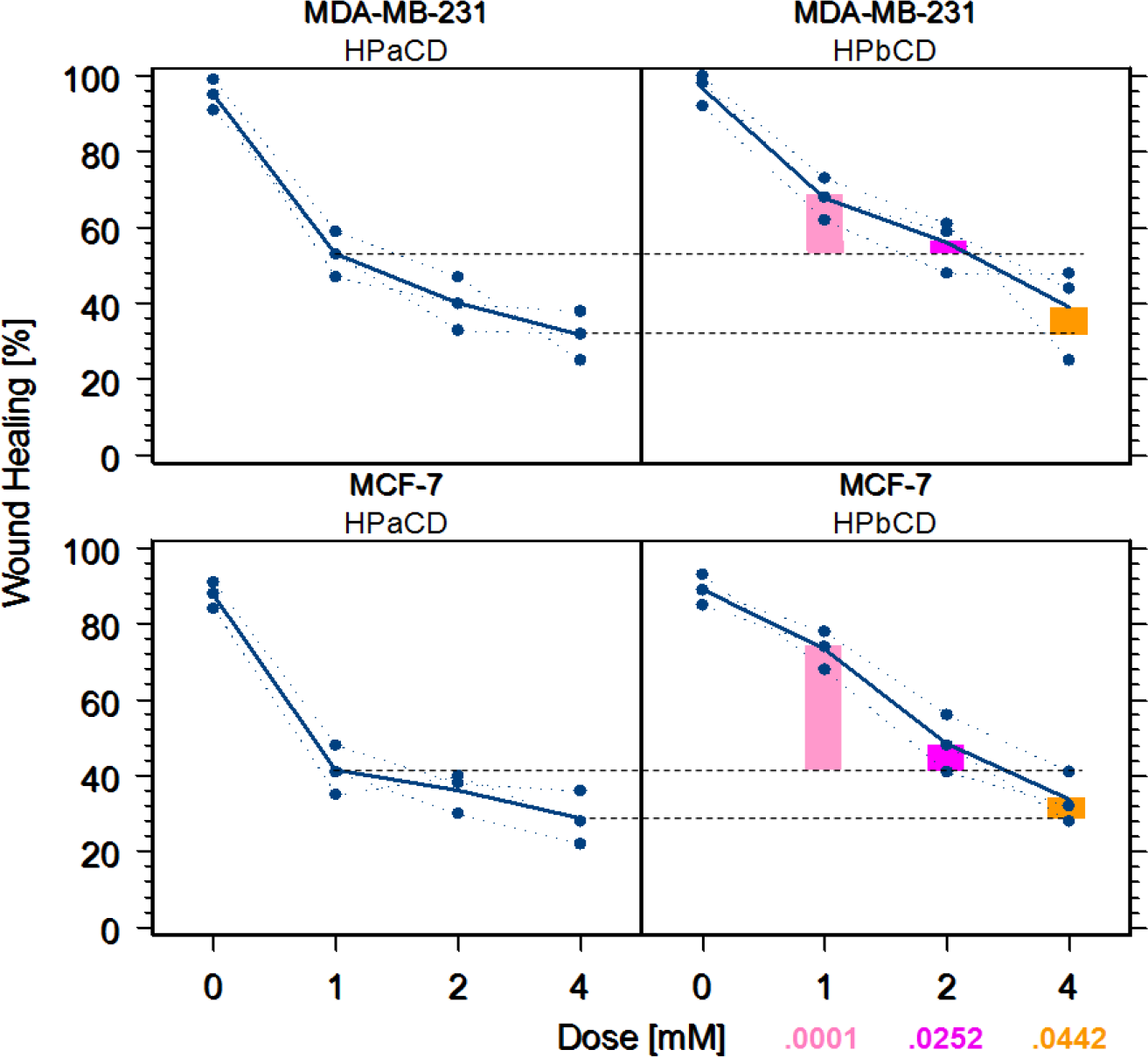
Wound healing by cyclodextrins in breast cancer cell lines. Cells were grown in triplicates for 12 h and incubated with either of the CDs for 2 h at the concentration indicated (0-4 mM), before a 0.9 mm wide gap was opened and cells were allowed to migrate into the “wound” for 12 h. HPβCD is more than 10× as toxic as HPαCD, which at <100 mM does not affect epithelial cell viability ^(Leroy-Lechat 1994; Roka 2015)^ Dashed horizontal line indicates inhibition of wound healing in HPαCD at 1and4mM respectively.

### ANOVA results

indep: HPαCD vs HPβCD (fixed)
block: MCF-7/MDA-MB-231 (fixed)
dep: %change in wound healing

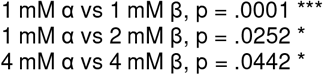

From Fig 6, 1 mM HPαCD is more effective than 2mM HPβCD against migration of ER^−^ and ER^+^ tumor cells (p= .0252) while more than 10× less toxic,^(Leroy-Lechat 1994)^ Hence, the effect previously seen with HPβCD is, in fact, likely the effect of it scavenging phospholipids, rather than cholesterol.

## Discussion

Our analysis confirmed previous GWAS, which pointed to receptor/AKT signaling and nuclear functions as critical components in breast cancer etiology. The present results from a reanalysis of data found previously inconclusive provides the first GWAS evidence for the contribution of EEC dysfunction and novel evidence for overstimulation of EEC in mesenchymal tumor cell migration and invasion. These findings, confirmed by an *in vitro* study on the activity of HPαCD vs HPβCD against breast cancer cell migration, suggest the novel hypothesis that reducing the influx of phospholipids, including PS, PC, and lysophosphatidylcholine (LPC), into the PI cycle via HPαCD could provide an urgently needed treatment option for women with breast cancer.

### Replication and complementation of previously identified genes

A previous analysis of the CGEM data reported only two genes, *FGFR2* and *BUB3*, as risk factors for breast cancer. The EPIC and PBCS data have been published only as part of three meta-analysis, which also included CGEM. Among ER^−^ cases, the first meta-analysis^(Siddiq 2012)^ confirmed two SNPs each in *BABAM1* (7.31) (a nuclear *BRCA1* complex component), *PTHLH* (12.8) (which regulates epithelial-mesenchymal interactions during the formation of mammary glands), and the ER *ESR1* (9.6). Our findings of *BMP7* (EPIC) and *BMPRT1B* (CGEM) are consistent with the previous finding of *PTHLH*, which forms a nuclear complex with *BMP4*. The second meta-analysis,^(Garcia-Closas 2013)^ pointed to the *PIK3C2B-MDM4* region (11.68), *LGR6* (7.85) (a GPCR), and *FTO* (7.40) (a regulator of nuclear mRNA splicing). Hence, ssGWAS in all three populations point to receptor/AKT signaling and nuclear processes, although the individual genes differ.

Three of the four EEC genes identified in previous ssGWAS;^(Michailidou 2013)^ were confirmed in muGWAS at aGWS/2 (CGEM: 2.56 / EPIC: 2.86 / PBCS: 2.57, S1 Table 1) in regions in LD (r^2^):^(Machiela 2015)^

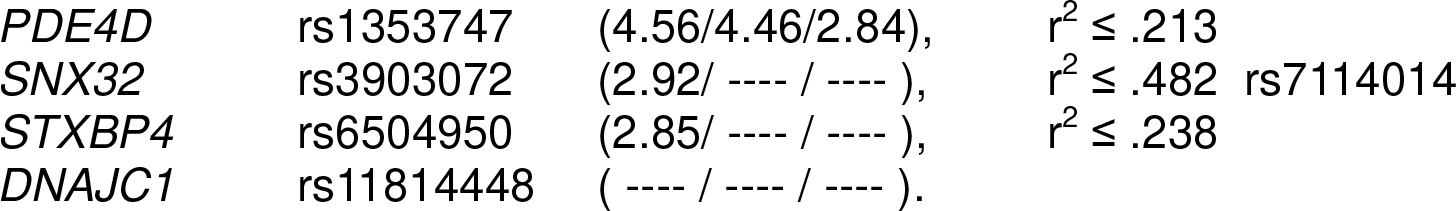

The EEC genes identified in here (with the exception of *AGPAT3/4, ASTN2*, and *EEA1*), have previously been shown to be associated with breast cancer in gene expression and functional studies (S1 Table 1).

A third meta-analysis ^(Hoffman 2017)^ based the above three and eleven other U4C data sets,^(Mechanic 2017)^ identified five novel breast cancer genes, three with nuclear function (*RCCD1, ANKL1, DHODH* ^(Mohamad Fairus 2017)^); *ACAP1* and *LRRC25* were hypothesized to be involved in cell proliferation (activating Arf6 protein) and inflammatory response (activating hematopoietic cells), respectively, ^(Hoffman 2017)^ In fact, both genes are can be related to EEC/PI in metastases: *ACAP1* (Fig 3, top right) regulates recycling of integrin β1 during cell migration ^(Li 2005)^; *LRRC25*, which regulates development of neutrophils needed for metastases,^(Coffelt 2016)^ carries a PI3K interaction motive.^(Liu 2017)^

A fourth single-SNP meta-analysis of 68 studies (including the three studies separately analysed here) with a total of 227,000 subjects ^(Michailidou 2017)^ identified “65 new breast cancer risk loci” to be “incorported into risk prediction models”, with “exocytosis” as the second-most significant “theme” (^(Michailidou 2017)^, Suppl Tab. 24).

### Computational biostatistics approach to genetic data

The analysis approach,^(Wittkowski 2013)^ used here integrates genetics concepts into the statistical method, rather than considering them during visual inspection of p-values calculated one SNP at a time and correlations among SNPs within an LDB. In particular, muGWAS avoids assumptions about the degree of dominance, reflects that both SNPs next to a disease locus should be in LD (unless they are separated by a recombination hotspot), increases resolution within LDBs (by distinguishing between members of the same tag sets being in a different order), integrates information from different disease loci within the same region (similar effects, compound heterozygosity), and draws on a measure of “information content” to prioritize results.

Screening for cis-epistatic regions (which may plausibly have evaded selective pressure) prioritizes biologically plausible results while de-emphasizing individual SNPs, which may be significant because of population selection biases, unless they cause exclusively late-onset phenotypes, such as age-related macular degeneration.^(Klein 2005)^ Avoiding strong model assumptions (additivity, independence) reduces model misspecification biases. Increasing the sample size, instead, does not guard against these biases, so that imposing a higher fixed GWS level in ssGWAS may, somewhat counterintuitively, favor “false positives” over biologically plausible cis-epistatic effects. The main limitation of u-statistics for multivariate data (conceived in the 1940s ^Hoeffding 1948)^) is that the amount of memory required became available only with 32-bit operating systems, in 2001, and computations became feasible only with the advent of GPU-enabled cloud computing.

To improve upon the conventional “overly conservative correction”^(Pearson 2008)^ of 7.3, a systematic analysis of GWA studies suggested lowering the GWS level to 7.0 (fixed),^(Panagiotou 2012)^ and then further by using study-specific empirical approaches.^(Aslibekyan 2013a)^ The empirical aGWS decision rule used here accounts for GWAS not being randomized, the absence of a traditional ‘null hypothesis’ in a heritable disease, differences in MAF causing the expected distributions in a QR plot to be convex, and tests in overlapping diplotypes being related.^(Wittkowski 2014)^

The combination of a method with higher specificity and a decision strategy with higher sensitivity increased the number of genes above the cut-off while ensuring that the vast majority of genes implicated was related to known pathways in breast cancer etiology, including dysregulation of EEC.

### Replication of findings across populations

Conventionally, a lower GWS level required for replication. At the aGWS/2 level, none of most significant ssGWAS results (CGEM: *FGFR2, BUB3, MMRN1*; EPIC: *CHD3, SOHLH2*; PBCS: *DOCK8*) was replicated in another population (S1 Table 2). Only three genes (*AGPAT3*, *MEGF11*, and *TRAPPC9*) were replicated in both of the other populations, but none for the same SNP. These results are consistent with common lack of replication in ssGWAS.^(Ioannidis 2013)^ With muGWAS, in contrast, many genes were replicated in at least one population and seven genes were replicated in both of the other populations (*FGFR2, TRPM3, AGPAT3, NCOR2*, *MEGF11, GPC6*, and *RGS3*), although not necessarily in the same intragenic region. Hence, analyses combining the data from several studies (often called “meta-analyses”, even when subject-level data is used) may result in some populations diluting the risk factors present in others.^(Ioannidis 2013)^

Our results are consistent with ssGWAS finding recent, highly penetrant mutations, which may differ across populations, while muGWAS has higher power for common cis-epistatic variations, which are more likely to be shared across populations. Even more likely to be shared are genes that carry different variations and different genes with similar contribution to the etiology,^(Aslibekyan 2013b)^ consistent with previous findings that breast cancer gene expression signatures have little overlap across populations.^(Haibe-Kains 2008)^

### Dysregulated EEC in breast cancer metastasis, angiogenesis, and progression

Genes involved in EEC (e.g., Rab GTPases) are aberrantly expressed in human cancers. ^(Mosesson 2008)^ Dysregulation of endocytosis-mediated recycling of oncoproteins (e.g., GF receptors and adhesion molecules, including integrins and annexins), can promote progression, migration, and invasion ^(Mosesson 2008; Maji 2016)^. Cell migration and invasion, which are promoted by EEC of integrins, are also essential features of angiogenesis.^(Demircioglu 2016)^ In addition, endocytic uptake of lipoproteins is critical for adaptation of cancer to its microenvironment.^(Menard 2016)^

Tumor-derived exosomes, 30-150 nm sized extracellular vesicles formed by dysregulated EEC, are critical mediators of intercellular communication between tumor cells and recipient stromal cells in both local and distant microenvironments.^(Costa-Silva 2015; Zhang 2015a)^ Several Rab proteins (Rab2b/5a/9a/27a/27b) are known to function in the selective packaging and production of exosomes in tumor cells (Fig 3, bottom left).^(Ostrowski 2010)^ Rab27a knockdown in highly metastatic melanoma cells significantly decreased exosome production, primary tumor growth, and metastasis,^(Peinado 2012)^ confirming the role of EEC in generating exosomes.

Dysregulated EEC alters not only exosome biogenesis (vesicular packaging and trafficking), but also the composition of exosomal cargos. Tumor-specific proteins, such as integrins were enriched in exosomes, transferred between cancer cells,^(Fedele 2015)^ and correlated with migration and invasion of recipient cells.^(Harris 2015; Keerthikumar 2015)^ Exosome uptake (involving endocytosis ^(Heusermann 2016)^) induces non-tumorigenic cells to develop cancer-related phenotypes and the uptake of exosomal integrins promotes migration of these tumor cells.^(Singh 2016)^

A recent study revealed that exosomal integrin expression patterns enriched in cancer-derived exosomes involve specific αβ combinations matched to target organs. Proteomic analysis revealed that the exosomal integrin αvβ5 binds to Kupffer cells that mediate liver metastasis, integrins α6β1 and α6β4 are associated with lung metastasis in breast cancer, while integrin β1 (which required for extravasation in metastases ^(Chen 2016)^) was not organ-specific. ^(Hoshino 2015)^.

Additionally, other tumor-specific exosomal proteins, such as annexins (calcium-dependent phospholipid-binding proteins known to regulate membrane trafficking and EEC), which are known to correlate with migration and invasion, are also packaged in cancer exosomes ^(Leca 2016; Maji 2016)^. Annexins are frequently overexpressed in breast, liver, prostate, and pancreatic cancers and participate in multiple functions in cancer, including angiogenesis, tumor migration and invasion.^(Mussunoor 2008)^ In breast cancer, exosomal annexin A2 promotes angiogenesis and vascularization via tissue plasminogen activator (tPA).^(Maji 2016)^ In pancreatic cancer, exosomal annexin A6 from cancer-associated fibroblasts contributes to tumor cell survival and invasion through annexin A6 / LDL receptor-related protein 1 / thrombospondin 1 complex formation.^(Leca 2016)^

In summary, EEC plays at least four roles in cancer development; spreading the cancer phenotype horizontally, preparing cancer cells for migration, preparing the distant microenvironment (all via preparation and transmission of exosomes containing integrins), and facilitating migration and invasion (via increasing EEC of integrins). In each case, both endo-and exocytosis are involved, either in donor and target cells or at trailing edge and advancing lamellipodium (Fig 2). Hence, down-regulating “de-railed endocytosis”^(Mosesson 2008)^ could have substantial synergistic effects.

### The PI cycle in Breast Cancer

Our findings of *PTENP1* (PBCS), *TNS1* (EPIC), and *SYNJ2* (CGEM) are consistent with known breast cancer mutations in PI3K/PTEN^(Varticovski 1991; Li 1997)^ and *SYNJ2*. That both PI(3,4,5)P_3_ and PI(3,4)P_2_ are required to achieve and sustain a malignancy, has been formulated as the “two PI hypothesis”^(Kerr 2011)^ Except for the known *PRCKQ*, which is regulated by phospholipids via the Pl(4,5)P_2_-PLC-DAG route, however, our analysis identified few genes along the *AKT/TSC/mTOR* pathway, which is controlled by the “two PIs”. Instead, our results point to EEC, in which virtually all PIs are involved. The closely related set of genes involved in recycling of DAG (*DGKQ*), influx of PC and PS (*ATP8B1, ATP8A1*), and influx of LPA and 1-acyl GPI (*AGPAT3, AGPAT4*) suggests the downregulation of circulating phospholipids as a novel strategy to reduce EEC.

LPA, a known promoter of cell migration and invasion in breast cancer,^Mills 2003; Wang 2016)^ is produced from LPC by autotaxin (ATX).^(Benesch 2016)^ While *ATX* mouse knockouts are embryonically lethal, mice that overexpress LPA or *ATX* develop spontaneous metastatic mammary tumors. A mechanism mediated by G-coupled LPA receptors may cause mesenchymal tumors via endocytosis upregulation involving β-arrestin2 ^(Alemayehu 2013)^ and Arf6.^(Hashimoto 2016)^

LPA and LPC in physiologic concentrations have been shown to strongly induce migration of rhabdomyosarcoma (RMS) cells and to be increased by irradiation and chemotherapy in bone marrow.^(Schneider 2014)^ The authors suggested the development of inhibitors of LPA/LPC signaling or “molecules that bind these bioactive lipids” after radio/chemotherapy. However, targeting a single among several redundant receptor/ligand complex may not be sufficiently effective to prevent metastases.^(Rataiczak 2016)^

Alkyl-LPCs, which compete with LPC, are in clinical use for treatment of cutaneous metastases in breast cancer, but have shown little activity (and substantial Gl side effects) in advanced metastatic breast cancer.^(Rios-Marco 2017)^ From the results presented here, this is consistent with reducing LPC being most effective while cells are still migrating.

As our results suggest, overall EEC upregulation may be caused by multiple variations affecting the PI cycle. Thus, reducing EEC by diminishing the overall phospholipid pool might be a more effective breast cancer treatment than blocking one or even two phosphotransferases, a strategy for which the highly robust PI cycle is designed to compensate. Given the ability of biologic systems to prioritize scarce resources, one would expect this effect to be stronger for tumor cells than for host cells whose functions are routinely prioritized when supplies are scarce. A related approach, substituted myo-inositol (Ml) analogues, had already been considered, but was found unlikely to be effective *in vivo*, because even physiological concentration of Ml antagonized the growth inhibitory activity of such analogues.^(Powis 1995)^

### βCDs are effective in cancer models of migration, invasion, and angiogenesis

A plethora of studies have investigated the effect of methyl-β-cyclodextrin (MβCD) *in vitro*. For instance, MβCD suppressed translocation of β1 integrin^(Huang 2006)^ and also invasion activity in three H7 Lewis lung cancer cell lines where highly metastatic cell lines had more β1 integrin.^(Zhang 2006)^. Breast and prostate cancer cell lines were more sensitive to MβCD-induced cell death than their normal counterparts.^(Li 2006)^ In particular, MβCD treatment induced a substantial decrease (40%) in activity of breast cancer resistance protein (*BCRP/ABCG2*),^(Storch 2007)^ which transports PS and PC analogues.^(Daleke 2007)^ In subsequent functional studies, MβCD inhibited spheroid migration and invasion of MDA-MB-241 and ZR751 breast cancer cells ^(Raghu 2010)^ and also endocytosis ^(Palaniyandi 2012)^ and migration ^(Guerra 2016b)^ of MCF7 breast cancer cells. MβCD was more toxic for invasive than for non-invasive urothelial cancer cells,^(Resnik 2015)^ and interfered with RTK-[PI2]-PI3K-[PI3]-AKT signaling in HeLa cells.^(Yamaguchi 2015)^ Finally, MβCD reduced breast cancer-induced osteoclast activity in RAW264.7 cells and osteoclastogenic gene expression in MCF-7 cells.^(Chowdhury 2017)^ Sulfated SβCD also inhibits epithelial cell migration and invasion, but not proliferation ^(Watson 2013)^ and prevents angiogenesis *ex vivo* in an rat aortic ring assay and an chick embryo collagen onplant assay.^(Watson 2013)^ The relevance of these *in vitro* findings was confirmed by several *in vivo* studies.

MβCD had higher concentration in tumor than in other cells (except kidney and liver involved in its clearance) and reduced tumor volume in mice xenografted with MCF-7 breast cancer or A2780 ovarian carcinoma cells at least as effectively and with less toxicity than doxycyclin,^(Grosse 1998)^ reduced the number of lung metastases in mice implanted with H7-O Lewis lung cancer cells,^(Zhang 2006)^ reduced invasiveness of melanoma,^(Fedida-Metula 2008)^ and inhibited growth of primary effusion lymphoma (PEL) in mice.^(Gotoh 2014)^ HPβCD was necessary in triple combination treatment for tumor regression in mice implanted with renal cancer cells.^(Yamaguchi 2015)^ and prolonged survival in leukemia mouse models.^(Yokoo 2015)^

βCDs have also seen effective in animal models of several other diseases known to involve endocytosis^(Coisne 2016)^: Alzheimer’s disease (APP),^(Yao 2012)^ Parkinson’s disease (α-synuclein),^(Bar-On 2006)^ and atherosclerosis (LDL),^(Montecucco 2015; Zimmer 2016)^. However, while HPβCD was well tolerated in most peripheral and central organ systems,^(Cronin 2015)^ it was shown to carry the risk of causing permanent hearing loss in mice,^(Crumling 2012)^ cats,^(Ward 2010; Vite 2015)^ and at least one human.^(Maarup 2015)^ Both intracochlear HPβCD and, in particular, MβCD were seen to be ototoxic in Guinea pigs.^(Lichtenhan 2017)^ This ototoxicity is believed to be due to depriving prestin (*SLC26A5*) in outer hair cells of cholesterol.^(Kamar 2012; Yamashita 2015; Takahashi 2016)^

### Migration and invasion in breast cancer involve cholesterol-unrelated processes

The role of phospholipids emerging from our results, however, suggests a different mechanism than scavenging of cholesterol. This mechanism is consistent with previously reported *in vivo* results: *CAV1* expression in breast cancer stroma increases tumor migration and invasion ^(Goetz 2011)^ and *CAV1* is required for invadopodia formation specifically by breast cancer cells, where *CAV1* knockdown cannot be rescued by cholesterol.^(Yamaguchi 2009)^ Growing MDA-MB-231 breast cancer cells in lipoprotein depleted medium resulted in an 85% decrease in cell migration.^(Antalis 2011)^ LPA activates the Arf6-based mesenchymal pathway for migration and invasion of renal cancer cells, which also originate from cells located within epithelial ductal structures. ^(Kamar 2012; Yamashita 2015; Hashimoto 2016; Takahasni 2016)^

Limiting the availability PIs would be particularly effective for PI(4)P and PI(4,5)P_2_ (each at <10%, see Fig 4) and, thus, would likely reduce endocytosis more than lysosomal degradation. In addition, cyclodextrins have been shown to exert their role in NPC treatment by activating rather than downregulating, Ca-dependent lysosomal exocytosis.^(Chen 2010)^

From the mechanism of βCD in NPC and elevated cholesterol levels seen in several cancers, including breast cancer,^(Yokoo 2015)^ βCDs were thought to reduce cancer growth by lowering cholesterol levels. Early evidence that this might not be the case emerged from the study of exosomes, which play a key role in development of breast cancer.^(Peinado 2011; Lowry 2015)^ Treatment of MDA-MB-231 breast cancer cells with MβCD inhibited the internalization of exosomes containing integrins,^(Hoshino 2015)^ but did so independently of cholesterol.^(Koumangoye 2011)^

### αCD scavenge phospholipids only, reducing AEs and increasing effectiveness

βCDs is widely believed to act through “cholesterol depletion”,^(Gotoh 2014; Badana 2016)^ yet βCDs also scavenges phospholipids.^(Ohtani 1989)^ From the genetics results, which suggest an overactiv PI cycle (Fig 4) for an age-related decrease of lysosomal function (Fig 3), the effect seen in breast cancer and some of the other diseases may be primarily through scavenging phospholipids. The cavity of αCDs is too small for cholesterol, but large enough for phospholipids.^(Rajnavolgyi 2014; Shityakov 2016)^ From the *in vitro* results validating the breast cancer hypothesis generated as part of the U4C challenge (Fig 6), αCDs may be more effective than βCD, yet without the risk of cholesterol-related ototoxicity.

Two types of “controls” have been used: repletion of cholesterol via βCDs “loaded” with cholesterol, and reduction of cholesterol production via statins. Repletion of cholesterol, however, also increases production of phospholipids by freeing acetyl-CoA, the precursor of both phospholipids and cholesterol,^(Shiratori 1994; Ridgway 1999; Lagace 2015)^ cholesterol replenishment restores sphingolipid decrease,^(Huang 2006)^ and statins also lower phospholipids.^(Snowden 2014)^ Hence, neither of these two strategies can “controls” against βCDs scavenging phospholipids, rather than cholesterols. Using αCD as a control, however, can answer this question and the above *in vitro* results suggest that equimolar αCDs are, in fact, at least twice as effective as βCDs, as one would expect if the effect of either CD is caused by its ability to scavenge phospholipids. Hence, our results suggest that many of the previous experiments with βCDs should be redone, this time using αCDs as a control.

αCD is generally recognized as safe (GRAS)^(FDA, GRN000155)^ and approved as an expedient for i.v. alprostadil.^(Loftsson 2010)^ Due to higher watersolubility, αCD has lower nephrotoxicity than βCD.^(Frank 1976)^ HP derivatives of αCD and βCD increase water solubility from 145 and 18.5, respectively to ≥500 g/L. In mice, the observed ototoxicity order of HPβCD >_[p<.002]_ HPγCD >_[p<.02]_ HPαCD [≈_[NS]_ vehicle] matches the reported order for hemolysis and toxicities in various cell types.^(Leroy-Lechat 1994; Davidson 2016)^ In humans, a single dose of up to 3 g/kg/d HPβCD and seven daily doses of 1 g/kg/d were reported to have no adverse effects.^(Gould 2005)^ In 5-yr old children treated for NPC, 800 mg/kg/d HPβCD i.v. for 12 months was well tolerated.^(Hastings 2009)^

### HPαCD as a potential novel treatment in breast cancer

Given significant redundancy pro-metastatic ligand-receptor complexes, the paradigm of targeting a single receptor-ligand complex has recently been challenged.^(Ratajczak 2016)^ Although targeting EEC is a promising therapeutic strategy to prevent and treat metastasis,^(Chew 2016)^ a therapeutic agent is yet to be determined. Our results suggest that metastases in breast cancer rely on upregulation of the highly robust PI cycle and various types of dysregulation along the complex EEC pathway, rather than a simple linear PI pathway. Hence targeting the PI cycle in its entirety may be more effective than targeting individual phosphatases or kinases, or specific genes along the EEC pathway. Cyclodextrins are attractive candidates for a polyvalent approach to treat breast cancer. By modulating several pathways involved in breast cancer, such as altering exosome production and packaging, and impede metastatic colonization, CDs are likely to confer greater protective effects than molecules that have single targets. The selectivity of the smaller αCDs to phospholipids would minimize side effects (e.g., ototoxicity) from βCDs also capturing cholesterol. Given that some CDs are already routinely used clinically, and their pharmacokinetic and toxicity profiles are well established, repeating previous encouraging animal studies of HPβCD, this time using HPαCD could lead rapidly to clinical efficacy trials. In women with TNBC, such trials could be completed within five years.

